# A novel defective recombinant porcine enterovirus G virus carrying a porcine torovirus papain-like cysteine protease gene and a putative anti-apoptosis gene in place of viral structural protein genes

**DOI:** 10.1101/510131

**Authors:** Ryo Imai, Makoto Nagai, Shoichi Sakaguchi, Tsuneyuki Masuda, Moegi Kuroda, Mami Oba, Yukie Katayama, Yuki Naoi, Shinobu Tsuchiaka, Tsutomu Omatsu, Hiroshi Yamazato, Shinji Makino, Tetsuya Mizutani

## Abstract

Enterovirus G (EV-G) belongs to the family of *Picornaviridae*. Two types of recombinant porcine EV-Gs carrying papain-like cysteine protease (PLCP) gene of porcine torovirus, a virus in *Coronaviridae,* are reported. Type 1 recombinant EV-Gs are detected in pig feces in Japan, USA, and Belgium and carry the PLPC gene at the junction site of 2C/3A genes, while PLPC gene replaces the viral structural genes in type 2 recombinant EV-G detected in pig feces in a Chinese farm. We identified a novel type 2 recombinant EV-G carrying the PLCP gene with flanking sequences in place of the viral structural genes in pig feces in Japan. The ~0.3 kb-long upstream flanking sequence had no sequence homology with any proteins deposited in GenBank, while the downstream ~0.9 kb-long flanking sequence included a domain having high amino acid sequence homology with a baculoviral inhibitor of apoptosis repeat superfamily. The pig feces, where the novel type 2 recombinant EV-G was detected, also carried type 1 recombinant EV-G. Although the phylogenetic analysis suggested that these two recombinant EV-Gs have independently evolved, type 1 recombinant EV-G might have served as a helper virus by providing viral structural proteins for dissemination of the type 2 recombinant EV-G.

## Introduction

Enterovirus G (EV-G) is an enveloped RNA virus, belonging to the family of *Picornaviridae*. More than 20 types of EV-G, known as EV-G1 to EV-G20, have been identified [1,2]. The viral genome is a single-stranded and positive-sense polarity and consisted of the 5’ untranslated regions (5’ UTR), one open reading frame (ORF), 3’ UTR and the 3’ end poly(A) sequence. In infected cells, one polyprotein is translated from the ORF and then processed to 4 structural proteins (VP1, VP2, VP3, VP4) and 7 nonstructural proteins (2Apro, 2B, 2C, 3A, 3B, 3Cpro, 3Dpol) via viral proteinases, 2A and 3CD.

By using a metagenomics approach, we have detected the genomes of EV-G1, 2, 3, 4, 6, 9, 10 and a new type of EV-Gs in feces of pigs with or without diarrhea in Japan [3]. Among them, 16 isolates of EV-G1 and one isolate of EV-G2 show an insertion of a papain-like cysteine protease (PLCP) gene from porcine torovirus at the 2C-3A junction sites [4] (Fig. 1A). We call them type 1 recombinant EV-G in this study. Our previous data demonstrate high prevalence of type 1 recombinant EV-G in the EV-G population. Type 1 recombinant EV-Gs have been discovered from feces of neonatal pigs showing clinical symptoms, such as diarrhea, in the US and Belgium [5,6,7], while type 1 recombinant EV-Gs identified in Japan are detected from normal as well as diarrhea pig feces [4]. In addition to type 1 recombinant EV-G, second type of recombinant EV-G (type 2 recombinant EV-G), which carried the PLCP gene in place of viral structural genes, has been identified in a Chinese pig farm [8].

**Fig. 1.**
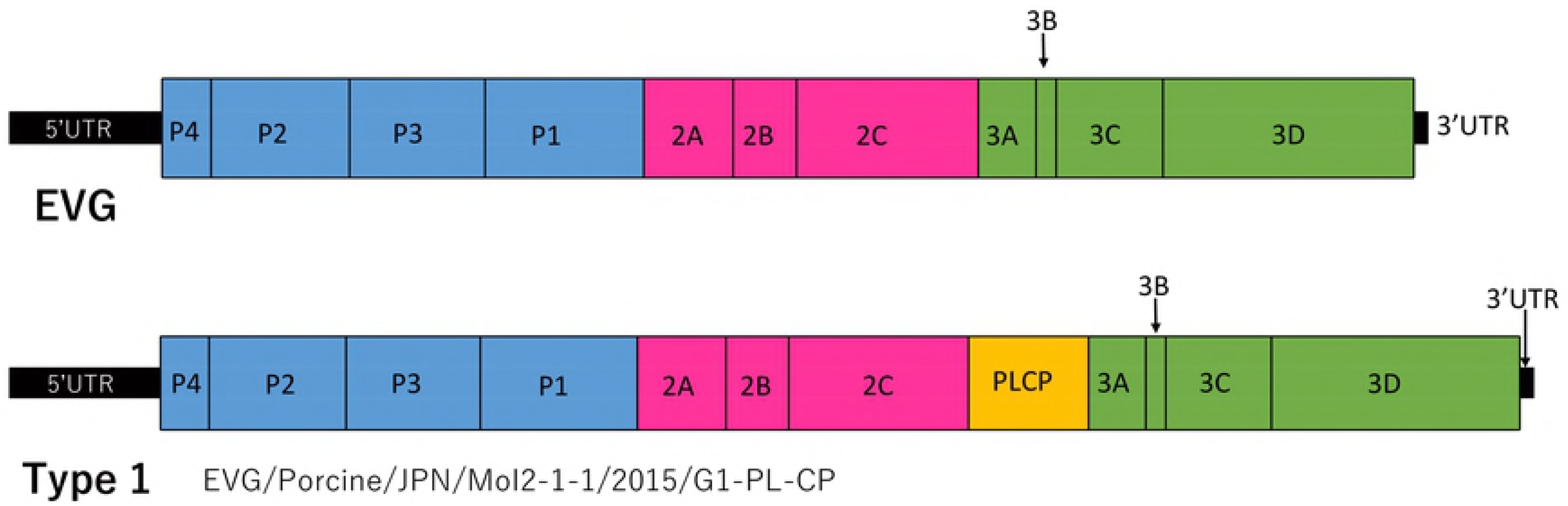
Schematic diagram of the genome organization of EV-G, type 1 (A) and type 2 recombinant EVGs (B). (A) Genome order of EV-G and type 1 recombinant EV-G was drawn from strain of EVG/Porcine/JPN/MoI2-1-1/2015/G1-PL-CP reported in our previous study [3]. (B) Genome order of a newly identified type 2 EV-G (EVG/Porcine/JPN/MoI2-1-2/2015/type 2) is shown in the top. The sequence of the newly identified type 2 EV-G was confirmed by overlapping PCR. The PCR products were electrophoresed using agarose gel (shown in left bottom panel). Lane 1, 31F and 1191R primers; lane 2, 930F and 1978R primers; lane 3, 1725F and 2760R primers; lane 4, 2500F and 3447R primers; lane 5, 3001F and 4188R primers; lane 6, 3921F and 4961R primers; lane 7, 4818F and 5852R primers; lane 8, 5619F and 6694R primers in Table 1. (C) Amino acid sequences at junction sites of unique region 1-PLCP-unique region 2-N-terminal truncated 2A of EVG/Porcine/JPN/MoI2-1-2/2015/type 2.

The PLCP gene is encoded in the ORF1 of the genome of torovirus, a member of family *Coronaviridae*, and the order *Nidovirales* [3,5-9]. The PLCP of nidoviruses serves as a protease to cleave viral gene 1 polyproteins to mature proteins [9]. It also has deubiquitinating and de-interferon stimulated gene removing enzyme (deISGylation) functions [5], which plays important roles in viral pathogenesis by acting as an innate immunity antagonist.

In the present study, we identified a novel type 2 recombinant EV-G in pig feces in Japan. In contrast to type 2 recombinant EV-G detected in the Chinese pig farm [8], the novel type 2 recombinant EV-G had an insertion of PLCP with flanking genes, one of which had a region showing high homology with a baculovirus gene having anti-apoptotic function.

## Materials and Methods

### Metagenomic analysis

Analysis of metagenomics data of RNAs of pig feces was reported previously [3]. We performed further analyses using the same metagenomics data and RNA samples.

### Long RT-PCR

RNAs were re-extracted from the pig feces using High Pure Viral Nucleic Acid Kit (Roche), and cDNA was synthesized with random primers using SuperScript Ⅲ First-Strand Synthesis System (Invitrogen), as describes previously [4]. Primers for PCR (Table 1) were designed to amplify the viral genome at approximately every 1 kb with overlap regions. PCR was performed by using GoTaq Master Mixes (Promega) with the following conditions: an initial denaturation at 95 °C for 2 min; followed by 35 cycles of 95 °C for 30 s, 55 °C for 30 s, and 72 °C for 1 min; and a final extension at 72°C for 5 min. We performed direct sequence analysis of the PCR products, which had been purified by agarose gel electrophoresis.

**Table 1.**
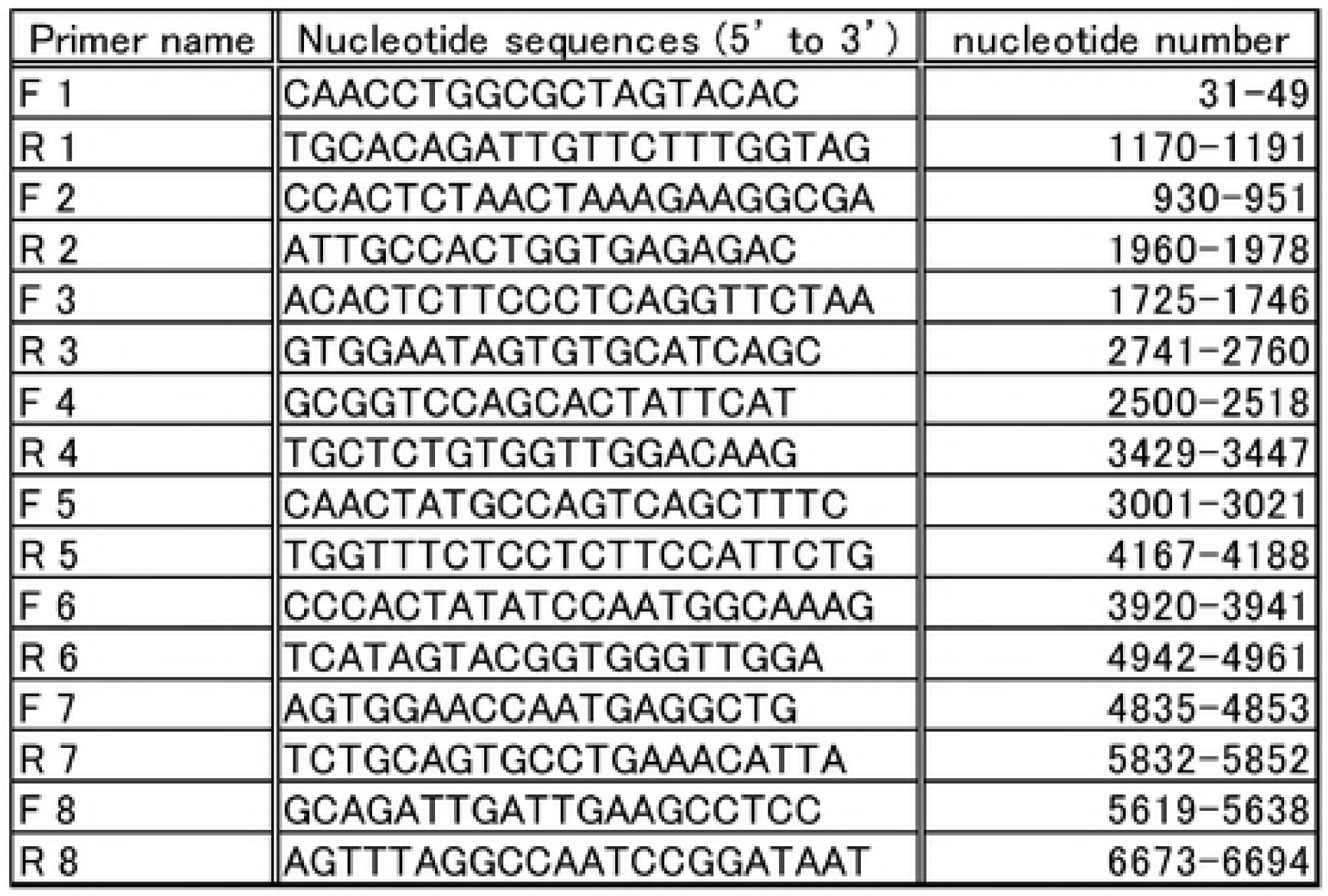
Primers for overlapping PCR

### Genome analysis

Nucleotide sequences were first subjected to a multiple sequence alignment using Clustal W [10] in MEGA 7 [11]. Evolutionary distance was estimated with the Kimura’s two parameter model [12]. Phylogenetic analyses were performed by the neighbor-joining method [13] using MEGA 7.

## Results

### Discovery of a novel type 2 recombinant EV-G in pig feces

Our previous metagenomics analyses of RNAs extracted from pig feces obtained from Japanese pig farms have identified various types of EV-Gs and type 1 recombinant EV-G [4]. Subsequent metagenomics analysis of the RNAs of the non-diarrheal pig feces obtained from a pig farm in Tottori prefecture, Japan, led to discovery of a novel type 2 recombinant EV-G genome (GenBank with accession number LC316774). This fecal sample also contains type 1 recombinant EV-G (EVG/Porcine/JPN/MoI2-1-1/2015/G1-Type 1) [4]. In contrast to the reported type 2 recombinant EV-G genome in China, which carries the PLCP gene of pig torovirus in place of viral structural genes [8], the newly identified type 2 recombinant EV-G (EVG/Porcine/JPN/MoI2-1-2/2015) carried the PLCP gene from pig torovirus plus ~0.3 kb-long upstream and ~0.9 kb-long downstream flanking sequences (Fig. 1B).

To confirm the genome sequence of the newly identified type 2 recombinant EV-G obtained by metagenomics analysis, we performed long RT-PCR to amplify the viral genome approximately every 1 kb with overlapping regions and subjected the PCR products for direct sequencing (Fig. 1B). These analyses revealed that the new type 2 recombinant EV-G genome was approximately 6.7 kb-long and had the following gene order: 5’ UTR - unique region 1 - PLCP – unique region 2 – 2A – 2B – 2C – 3A – 3B – 3C – 3D – 3’ UTR (Fig. 1B). The new type 2 recombinant EV-G genome carried a single ORF, suggesting that translation of the viral polyprotein starts at the translational start codon (ATG) at the 5’ end of the unique region 1 and ends at the stop codon (TAA) of 3D gene. In addition to the replacement of the structural genes with the PLCP gene carrying flanking regions, the newly identified type 2 recombinant EV-G lacked 11 amino acids at the N-terminal of 2A protein. We noted that the same 11 amino acids are also deleted in type 2 recombinant EV-G detected in the pig farm in China [8]. In contrast, the N-terminal truncation of the 2A proteinase gene did not occur in type 1 recombinant and non-recombinant EV-Gs (data not shown), implying that the truncated 2A may be a specific characteristic feature among type 2 recombinant EV-Gs. Furthermore, type 2 recombinant EV-Gs lacked GxCG motif in 2A, which is considered to form part of the active site of the protease and is conserved among enteroviruses [14]. In contrast all type 1 recombinant EV-Gs carry this motif in 2A (data not shown). Fig. 1C showed amino acid sequences at the junction sites of the unique region 1, PLCP gene, unique region 2, and the 2A lacking the N-terminal 11 amino acids [14].

### Unique regions of type 2 recombinant EV-G

An internal region (amino acids 180-240) of the 304 amino acid-long unique region 2 of the newly identified type 2 recombinant EV-G showed high amino acid homology with genes of baculoviral inhibitor of apoptosis repeat (BIR) superfamily (Fig. 2). In contrast, unique region 1 and other regions in unique region 2 showed no significant amino acid homologies with any known proteins.

**Fig. 2.** Amino acid alignment of unique region 2 and BIR family. Amino acid sequence of amino acids at 180-240 of unique region 2 of the newly identified type 2 EV-G is shown in top and is aligned with other BIR family protein. moi212uniq2 represents the newly identified type 2 EV-G. The accession number of protein sequences from GenBank used in this figure is indicated as XP_004837139.1: baculoviral IAP repeat-containing protein 8-like of Heterocephalus glaber, XP_020040963.1: E3 ubiquitin-protein ligase XIAP of Castor canadensis, XP_021484438.1: E3 ubiquitin-protein ligase XIAP of Meriones unguiculatus, XP_023055645.1: E3 ubiquitin-protein ligase XIAP of Piliocolobus tephrosceles, XP_023590042.1: E3 ubiquitin-protein ligase XIAP isoform X2 of Trichechus manatus latirostris, XP_020935024.1: E3 ubiquitin-protein ligase XIAP isoform X1 of Sus scrofa, XP_004480589.1: E3 ubiquitin-protein ligase XIAP of Dasypus novemcinctus, XP_024602944.1: E3 ubiquitin-protein ligase XIAP isoform X2 of Neophocaena asiaeorientalis asiaeorientalis, XP_022417552.1: E3 ubiquitin-protein ligase XIAP isoform X1 of Delphinapterus leucas, NP_001164796.1: E3 ubiquitin-protein ligase XIAP of Oryctolagus cuniculus, XP_007111017.1: E3 ubiquitin-protein ligase XIAP isoform X1 of Physeter catodon, XP_020140715.1: E3 ubiquitin-protein ligase XIAP isoform X3 of Microcebus murinus, EPY73910.1: Baculoviral IAP repeat-containing protein 4 of Camelus ferus, EGW11922.1: Baculoviral IAP repeat-containing protein 4 of Cricetulus griseus, NP_001271387.1: E3 ubiquitin-protein ligase XIAP of Canis lupus familiaris, XP_013003432.1: E3 ubiquitin-protein ligase XIAP of Cavia porcellus, XP_008068909.1: E3 ubiquitin-protein ligase XIAP of Carlito syrichta, AAB58376.1: X-linked inhibitor of apoptosis of Mus musculus, OWJ99336.1: XIAP of Cervus elaphus hippelaphus, XP_023363885.1: E3 ubiquitin-protein ligase XIAP of Otolemur garnettii.

### Phylogenetic analysis of type 2 recombinant EV-G

Phylogenetic analysis showed that PLCP, 2A, 2B, 2C, 3C and 3D genes of the newly identified type 2 EV-G made different clusters from known non-recombination EV-Gs and type 1 recombinant EV-Gs (Fig. 3A to 3F), including type 1 recombinant EV-G, which co-existed with the new type 2 recombinant type 2 in the same feces; type 1 shown in blue and type 2 shown in red were found in the same feces used in this study. These data suggest that the type 2 recombinant EV-G was not directly derived from known type 1 recombinant EV-Gs or non-recombinant EV-Gs. We also noted that the new type 2 recombinant EV-G had the 247 nt-long 3’ UTR (data not shown), which was substantially longer than 40 to 165 nt-long 3’ UTR of most of picornaviruses, supporting the notion that the new type 2 recombinant EV-G has evolved independently from type 1 recombinant EV-Gs or non-recombinant EV-Gs.

**Fig. 3.** Phylogenetic trees of EVG/Porcine/JPN/MoI2-1-2/2015/type 2 with enterovirus G strains from GenBank database based on nucleotide sequences of (A) PLCP, (B) 2A, (C) 2B, (D) 2C, (E) 3C, and (F) 3D coding regions. The trees were constructed using Neighbor-joining method in MEGA 7.0.14 and bootstrap test (n-1000). The genetic distance was calculated using Kimura’s two parameter model. The scale bar indicates nucleotide substitutions per site.

## Discussion

In the present study, we reported the genome of a novel type 2 recombinant EV-G from pig feces in Japan. The newly identified type 2 recombinant EV-G lacked the genes for structural proteins, while it carried most of the genes encoding viral nonstructural proteins. Accordingly, this defective recombinant EV-G required helper virus, which should provide viral structural proteins for dissemination, and underwent RNA replication in the absence of helper virus. Because the 1 recombinant EV-G was detected in the same feces sample [3] as the new type 2 recombinant EV-G, this type 1 recombinant EV-G, which belongs to different subtype, might have served as the helper virus.

Some RNA viruses acquire a new function, e.g., inhibition of the host immune functions, by gaining a new gene via RNA-RNA recombination [14,15]. Enteroviruses are genetically and antigenically highly variable due to recombination within as well as between serotypes [16,17]. Poliovirus and coxsackievirus undergo RNA recombination, with higher efficiency for non-homologous RNA recombination than homologous RNA recombination, in cell cultures, suggesting that non-homologous RNA recombination may be a transient and intermediate step for the generation and selection of the fittest homologous recombinants [18]. Identification of two different types of recombinant EV-G implied that non-homologous recombination between picornavirus RNA and non-picornavirus genes drives evolution of picornaviruses.

Although both type 2 and type 1 recombinant EV-Gs carry the PLCP gene derived from torovirus, location of the PLCP gene in the genome of the two recombinant EV-Gs differ (see Fig. 1A and 1B). As PLCP is known to have deISGylation activity [4], retention of torovirus PLCP gene in the two different recombinant EV-G types imply that the PLCP gene may suppress host innate immune functions, facilitating survival of these recombinant EV-Gs. The cluster of PLCP was different between recombinant EV-G type 1 and the newly identified recombinant EV-G type 2 (Fig. 3A), suggesting that these two different types of EV-Gs acquired the PLCP by RNA recombination from different subtypes of toroviruses.

Unlike most of recombinant EV-Gs, the newly identified type 2 recombinant EV-G carried the unique region 2, which had a domain showing extensive amino acid homology with the BIR superfamily. The baculoviral inhibitor of apoptosis (IAP) protein facilitates viral replication by preventing apoptosis [19,20]. Generally, IAP proteins contain one to three BIR domains, which play an important role in the anti-apoptotic function. IAPs have a RING domain, which relates E3 ubiquitin ligase activity for ubiquitination of target proteins degradation, at the C-terminal region [21,22,23]. Possibly, the BIR-like domain and the PLCP gene inhibited apoptosis and host innate immune function, respectively, leading to efficient replication of the newly identified type 2 EV-G.

One unanswered question in this study was whether the protein region translated from the non-picornaviral genes of the newly identified type 2 recombinant EV-G undergoes protein processing. Coronavirus PLCP cleaves the viral gene 1 polyproteins through recognition of a LXGG motif [8], while PLCP of arterivirus, another nidovirus, recognizes sequence LIGG, TTGG or PSGG [8]. However, cleavage specificity of torovirus PLCP has not need identified. Accordingly, it is unclear whether the PLCP in the newly identified type 2 recombinant EV-G cleaves any regions in the protein region translated from the inserted foreign genes. Enterovirus 2A and 3CD cleave Y/G pair and Q/G pair, respectively [24], while these pairs were absent at junctions at unique region 1/PLCP, PLCP/unique region 2 and unique region2/truncated 2A as well as within the putative polyprotein translated from the non-picornavirus sequences. Moreover, the absence of GxCG motif conserved in chymotrypsin-like protease in 2A of type 2 recombinant EV-Gs suggests that the 2A proteinase function of the type 2 recombinant EV-G 2A was most probably defective. As enterovirus 2A induces cleavage in host proteins [24], absence of biologically active 2A potentially affect host environment, including translational status, and possibly affect viral gene expression. Taken together, it is unlikely that the protein region translated from the inserted foreign genes undergo processing by the truncated 2A and 3CD.If the torovirus PLCP does not induce a cleavage(s) into the polyprotein of the newly identified type 2 recombinant EV-G, a protein consisted of unique region 1, PLPC, unique region 2 and N-terminal truncated 2A, might have been accumulated in cells infected with the newly identified type 2 recombinant EV-G. If so, testing the deISGylation function and anti-apoptotic function of this putative protein would provide a clue as to a possible reason(s) for retention of these nonpicornavirus sequences in the type 2 recombinant EV-G. Alternatively, the putative polyprotein with the torovirus PLCP and the BIR domain may have another biological function that is important for replication of the newly identified type 2 recombinant EV-G.

## Conclusions

A novel type of recombinant enterovirus G (type 2 recombinant EV-G) was discovered in pig feces in Japan. This type 2 recombinant EV-G carried the PLCP torovirus gene with an upstream flaking gene of unknown function and a downstream flanking gene having putative anti-apoptosis function, in place of the viral structural proteins. The phylogenetic analysis showed this type 2 recombinant EV-G belonged to be a different cluster from the cluster of type 1 recombinant, all of which were detected in the same sample.

## Acknowledgements

This study is supported by funding including JSPS KAKENHI 15K07718, and operating cost from Global Innovation Research of Tokyo University of Agriculture and Technology. We thank Dr. Jayne K. Makino for proofreading of the manuscript.

## Author Contributions

**Conceptualization**: TM MN. SM.

**Data Curation**: RI. SS. MK. MO.

**Formal Analysis**: MN. YN. ST.

**Funding Acquisition**: TM NM.

**Investigation**: RI. SS. MK. MO. YK. YN. ST. TO. HY.

**Methodology**: TM. RI MN.

**Project Administration**: TM MN.

**Resources**: TM. SS. SM.

**Software**: SS. YN.

**Supervision**: SM TM NM.

**Validation**: RI. MN.

**Visualization**: HY. TO.

**Writing – Original Draft**: RI. TM. SM.

**Writing – Review & Editing**: SS SM TM MN.

